# Forelimb unloading impairs glenohumeral muscle development in growing rats

**DOI:** 10.1101/2020.02.26.967273

**Authors:** Sophia K. Tushak, Margaret K. Tamburro, Emily B. Fawcett, Lauren E. Merritt LE, Katherine R. Saul, Jacqueline H. Cole

## Abstract

Proper joint loading is essential for healthy musculoskeletal development. Many pediatric neuromuscular disorders cause irreversible muscle impairments resulting from both physiological changes and mechanical unloading of the joint. While previous studies have examined the effects of hindlimb unloading on musculoskeletal development in the lower limb, none have examined solely forelimb unloading. Thus, a large deficit in knowledge of the effect of upper limb unloading exists and must be addressed in order to better understand how the glenohumeral joint adapts during development. Two forelimb unloading models were developed to study the effects of varying degrees of unloading on the glenohumeral joint in growing rats: forelimb suspension (n=6, intervention 21 days post-natal) with complete unloading of both limbs via a novel suspension system and forearm amputation (n=8, intervention 3-6 days post-natal) with decreased loading and limb use in one limb after below-elbow amputation. After 8 weeks of unloading, changes in muscle architecture and composition were examined in ten muscles surrounding the shoulder. Results were compared to control rats from a previous study (n=8). Both methods of altered loading significantly affected muscle mass, sarcomere length, and optimal muscle length compared to control rats, with the biceps long head and triceps long head observing the most marked differences. Forearm amputation also significantly affected muscle mass, sarcomere length, and optimal muscle length in the affected limb relative to the contralateral limb. Muscle composition, assessed by collagen content, remained unchanged in all groups. This study demonstrated that forearm amputation, which was administered closer to birth, had greater effects on muscle than forelimb suspension, which was administered a few weeks later than amputation.

## Introduction

Mechanical loading is critically important for healthy musculoskeletal development^1,2^ and maintenance^3,4^. In adult murine models, unloading via hindlimb suspension, microgravity during spaceflight, and muscle paralysis causes changes in muscle architecture. For example, unloading in adult murine animals caused substantial reductions of 41-66% in skeletal muscle size, mass, and strength^6,8,9^, as well as up to 13% longer sarcomere lengths^8^. However, muscle composition measured by collagen content was unaffected by unloading in adult rodents^7,11^. In growing animals, unloading is particularly impactful, causing irreversible musculoskeletal changes^5,12,56^, including altered joint morphology^12^, which influences surrounding muscle, and decreased muscle mass by over 3 to 5-fold^5,56^. However, the effects of unloading on other muscle architecture metrics (e.g., sarcomere length, optimal muscle length) and muscle composition (e.g., collagen content) have not previously been examined in growing animals.

Unloading models have traditionally focused on the hindlimbs^15^, resulting in limited understanding of the specific contributions of forelimb unloading to changes in muscle of glenohumeral joint, particularly during development. With a combined incidence of more than 5 per 1,000 live births^16–19^, pathologies affecting the developing muscles surrounding the glenohumeral joint (e.g., brachial plexus birth injury^20–22^, congenital muscular dystrophy^23–25^, cerebral palsy^26–28^, and congenital myasthenia gravis^29–31^) have substantial implications for the effects of altered forelimb loading during development. However, isolating the role that altered loading plays in these conditions is challenging, since, for example, nerve injury also directly contributes to detrimental muscle^12,13^ changes. Hence, the independent contributions of altered mechanical loading and nerve injury or other congenital changes to musculoskeletal development following neuromuscular disorders and injuries are difficult to elucidate. Understanding the contributions of unloading to these pathologies is an important step in determining which changes result directly from injury or disease and which are a functional consequence of altered joint loading, which may aid treatment development to restore muscle function.

Because of anatomical similarities to human shoulders^32,33^ and rapid development to skeletal maturity^34^, murine models are often used to study these injuries and diseases^20–31^. While murine hindlimb unloading^35^ and partial unloading paradigms exist^36^, to date no study has implemented a partial or complete forelimb unloading paradigm that targets the glenohumeral joint during neonatal development. A previous mouse model of bipedal locomotion training to improve function following spinal cord injury involved forelimb unloading, although these effects were not evaluated, and the animal was placed in an unnaturally upright posture, making this model unsuitable for studying long-term forelimb unloading^37^. We implemented two novel murine models to simulate variable degrees of mechanical unloading in the glenohumeral joint that occurs with this array of neuromuscular diseases and injuries: forelimb suspension and forearm amputation. Our objective was to determine the effect of these forelimb unloading models on the postnatal development of muscles surrounding the glenohumeral joint, in particular muscle architecture and composition.

## Methods

All procedures were approved by the Institutional Animal Care and Use Committee at North Carolina State University prior to the start of the study. Male and female Sprague Dawley rats (Charles River Laboratories, Wilmington, MA) were subjected to forelimb unloading using one of two different methods (Fig. 1): forelimb suspension (n = 6) or forearm amputation (n = 8). Results from the unloading groups were compared to results from a control group (n = 8).

**Figure 1.**
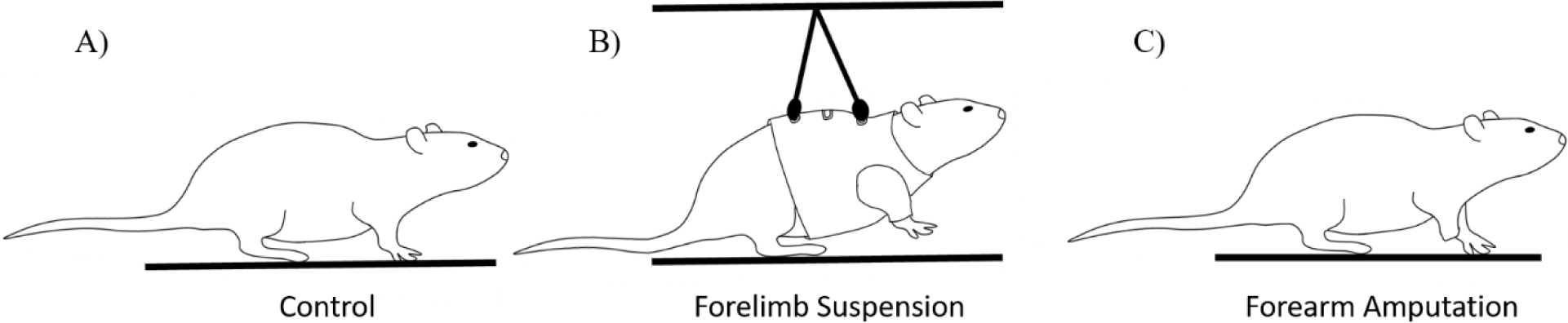
The study design included a A) control group (both forelimbs unaffected by unloading; right forelimb examined) and two unloading paradigms, B) forelimb suspension (both forelimbs affected) and C) forearm amputation (right forelimb affected, left forelimb unaffected).

### Forelimb Suspension

Six Sprague Dawley rats (2 female, 4 male) from three litters were exposed to forelimb unloading promptly after weaning at 3 weeks of age. Rats were placed in fitted harnesses, connected to a custom suspension system, and subjected to a six-week period of continuous unloading in both forelimbs (Fig. 2). Details of our suspension system were described previously^38^. The rats experienced a 12-hour light/12-hour dark cycle. Rat chow (Purina, Woodstock, Ontario, Canada) and HydroGel® (ClearH2O®, Inc., Westbrook, ME) were offered *ad libitum*. HydroGel® was used instead of water, because the typical water bottle interfered with the suspension system. At 9 weeks of age and after six weeks of loading, the rats were euthanized with CO_2_ inhalation followed by bilateral thoracotomy.

**Figure 2.**
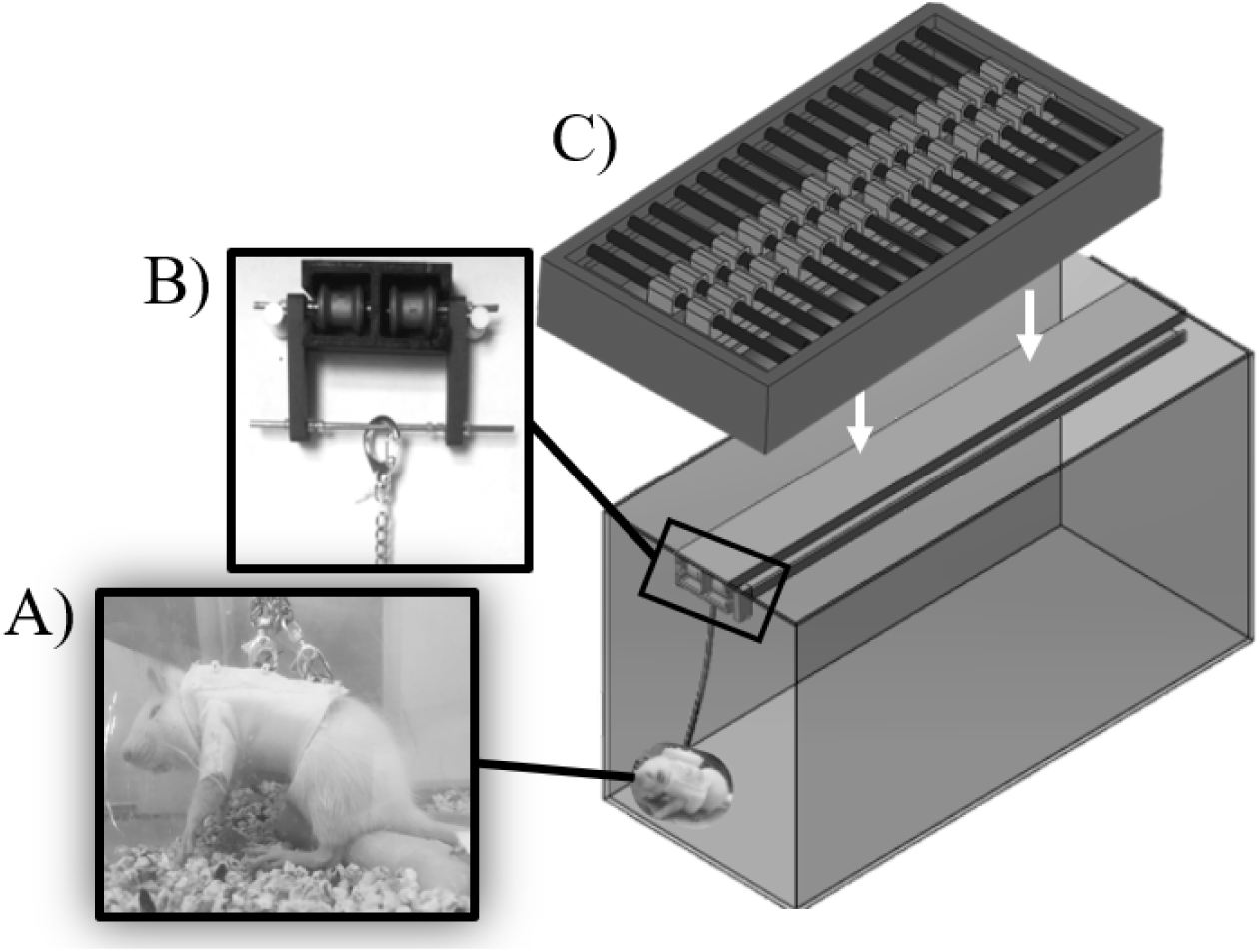
Custom forelimb suspension apparatus. A) Commercial harness sized to the growing rat was tethered at two points using swivel hooks and adjustable chain and attached to the B) 3D printed I-beam track system with low-friction wheels. C) Sixteen wooden dowels were inserted into loops printed atop the track system to secure it within the lid.

### Forearm Amputation

Eight Sprague Dawley rat pups (2 female, 6 male) from the same three litters as the forelimb suspension group received forearm amputations at 3-6 days of age. Rat pups were anesthetized with isoflurane, and right forearms were amputated through elbow disarticulation using aseptic technique, with the contralateral forelimbs remaining intact. The wound was irrigated and closed with tissue adhesive and suture. To minimize pain, rats received a local anesthetic (bupivacaine) at the incision site during surgery, one dose of buprenorphine (0.01 mg/kg) and carprofen (5 mg/kg) immediately after surgery, and a course of carprofen once per day for five days after surgery. Upon recovery from anesthesia, rat pups were returned to their dams and regularly monitored for signs of acceptance. Rat pups were weaned from their dams at 3 weeks of age and housed three per cage in the same room with the same accommodations as the forelimb suspension group, except they were given a typical water bottle. At 8 weeks of age, the rats were euthanized with CO_2_ inhalation followed by bilateral thoracotomy.

### Control

Control comparison data were obtained from previously assessed rats that underwent a sham surgery^50^. In that study, 8 Sprague Dawley rat pups (3 female, 5 male) from three litters received sham surgeries at 3-6 days of age that exposed the brachial plexus nerve bundle through the right pectoralis major, but no subsequent nerve injury was administered, and the contralateral forelimbs were kept intact. The wound was irrigated and closed with tissue adhesive. To minimize pain, one dose each of buprenorphine and carprofen was administered immediately following surgery. Rats received the same post-surgical care as the forearm amputation group. At 8 weeks of age, the rats were euthanized with CO_2_ inhalation followed by bilateral thoracotomy. For the control group, the left forelimb – which did not undergo surgery – was considered unaffected and used for comparison to the unloading groups.

### Muscle Dissection

Following euthanasia, the upper body was harvested using a guillotine to remove both the head and lower body. The torso was then fixed in 10% neutral buffered formalin for two days and stored in 70% ethanol at 4°C until muscle dissection. In 11 rats (5 control, 3 suspension, 3 amputation), 10 muscles surrounding the shoulder and upper forelimb were dissected bilaterally and stored in 70% ethanol at 4°C until architecture analysis: pectoralis major, acromiodeltoid, spinodeltoid, biceps long head, biceps short head, subscapularis, supraspinatus, infraspinatus, teres major, and triceps long head^39^. In the remaining 11 rats (3 control, 3 suspension, 5 amputation), four muscles (biceps long head, biceps short head, upper and lower subscapularis) were harvested bilaterally for composition analysis. The proximal end of each muscle was embedded in optimum cutting temperature compound and set in 2-methylbutane cooled by liquid nitrogen, and the entire muscle was then snap frozen and stored at −80°C until sectioning.

### Optimal Muscle Length

Muscle mass and muscle length were measured for the muscles stored at 4°C. After blotting excess ethanol, muscles were weighed on a digital scale (resolution of 0.01 g). For each muscle, 9 muscle fibers were extracted, 3 each from the proximal, middle, and distal regions of the muscle. Sarcomere lengths were measured via a 5.0-mW HeNe laser with a wavelength of 633 nm (Thorlabs, Newton, NJ) using an established laser diffraction method^26^. All muscle lengths and distances between each diffraction band were measured using digital calipers (resolution of 0.01 mm). The 9 sarcomere measurements were averaged to find the mean sarcomere length for each muscle. To determine the excursion capacity of the muscles and account for possible stretch in the fixed muscle as indicated by sarcomere length, optimal muscle length was calculated^40^:

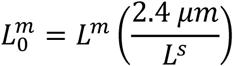

where *L*^*m*^ is muscle length and *L*^*s*^ is sarcomere length. The optimal sarcomere length corresponded to that of rat skeletal muscle (2.4 µm).

### Muscle Fibrosis

In muscles stored at −80°C, three transverse cryosections with a thickness of 10 µm were obtained from each muscle (Cryotome FSE Cryostat, Thermo Scientific, Halethorpe, MD), mounted to a silanized slide, and stored at −80°C prior to staining. Muscle sections were stained with Masson”s trichrome (American MasterTech, Lodi, CA) to identify collagen I deposition, a measure of fibrosis and muscle stiffening, and imaged at 20X magnification with light microscopy (EVOS^®^ FL Cell Imaging System, Thermo Scientific, Halethorpe, MD) In three sections per muscle, collagen content was calculated as the ratio of collagen area to muscle tissue area using a custom image processing protocol (MATLAB^®^, The MathWorks, Inc., Natick, MA).

### Statistical Analyses

To verify whether side-to-side differences were insignificant for the forelimb suspension and control groups (as expected) and to identify whether differences existed in muscle metrics between the affected and unaffected forelimbs for the amputation group, paired t-tests were used. Muscle architecture (mass, sarcomere length, optimal muscle length) and composition (collagen content) metrics were compared across the three groups (control, forelimb suspension, forearm amputation) using one-way ANOVA with Tukey’s post-hoc tests. For the group comparisons, data from only one forelimb was used: right for both unloading groups and left (unoperated) for the control group. All analyses were performed in RStudio Cloud (v. alpha, The R Foundation for Statistical Computing, Vienna, Austria) with a significance level of α = 0.05.

## Results

### Side Differences Within Each Group

In the control and forelimb suspension groups, no significant side-to-side differences were found for any of the metrics, apart from the pectoralis major muscle mass in the control group, which was lower in the sham limb (right) compared to the unimpaired limb (left) as previously reported^39^. This was expected, because the sham surgery involved a transverse infraclavicular incision through the pectoralis major to expose the brachial plexus. In the forearm amputation group, muscle mass was an average of 37.7% lower for muscles in the right (affected) limb compared to the left (unaffected) limb (Table 1). Affected limb muscle mass was significantly lower than unaffected for acromiodeltoid (18.1 ± 1.7%, p = 0.0153), biceps long head (54.9 ± 7.9%, p = 0.0136), and triceps long head (56.8 ± 6.0%, p = 0.0136).

**Table 1.** Muscle mass, sarcomere length, and optimal muscle length for the control (left forelimb), forelimb suspension (right forelimb), and forearm amputation (both forelimbs) groups. Mean ± standard deviation.

|  | control |  |  | forelimb suspension |  |  |
| --- | --- | --- | --- | --- | --- | --- |
|  | left (unaffected) |  |  | right (affected) |  |  |
| | mass<br>(g) | sarcomere length<br>(μm) | optimal length<br>(mm) | mass<br>(g) | sarcomere length<br>(μm) | optimal length<br>(mm) |
| pectoralis major | $0.34 \pm 0.08^a$ | $2.33 \pm 0.27$ | $19.4 \pm 2.1$ | $0.23 \pm 0.03$ | $1.92 \pm 0.08$ | $22.2 \pm 3.3$ |
| acromiodeltoid | $0.14 \pm 0.03$ | $2.49 \pm 0.14$ | $11.3 \pm 1.4$ | $0.10 \pm 0.03$ | $2.15 \pm 0.10$ | $14.2 \pm 1.2$ |
| spinodeltoid | $0.14 \pm 0.03$ | $1.90 \pm 0.04$ | $20.1 \pm 1.8$ | $0.12 \pm 0.05$ | $2.06 \pm 0.04$ | $25.0 \pm 2.4$ |
| biceps long head | $0.10 \pm 0.02$ | $2.26 \pm 0.11$ | $15.2 \pm 2.0$ | $0.10 \pm 0.02$ | $2.07 \pm 0.04$ | $16.8 \pm 1.3$ |
| biceps short head | $0.01 \pm 0.01$ | $2.09 \pm 0.21$ | $17.3 \pm 2.4$ | $0.04 \pm 0.02$ | $1.95 \pm 0.06$ | $21.1 \pm 1.4$ |
| subscapularis | $0.27 \pm 0.05$ | $2.34 \pm 0.10$ | $18.2 \pm 3.0$ | $0.19 \pm 0.06$ | $2.03 \pm 0.15$ | $21.6 \pm 0.6$ |
| supraspinatus | $0.21 \pm 0.06$ | $2.59 \pm 0.13$ | $20.2 \pm 2.2$ | $0.20 \pm 0.05$ | $2.21 \pm 0.14$ | $23.7 \pm 1.0$ |
| infraspinatus | $0.22 \pm 0.05$ | $2.37 \pm 0.10$ | $22.1 \pm 2.6$ | $0.19 \pm 0.02$ | $2.22 \pm 0.05$ | $24.0 \pm 2.0$ |
| teres major | $0.21 \pm 0.05$ | $1.72 \pm 0.04$ | $24.5 \pm 2.8$ | $0.21 \pm 0.05$ | $1.98 \pm 0.09$ | $25.0 \pm 0.9$ |
| triceps long head | $0.73 \pm 0.16$ | $1.98 \pm 0.13$ | $24.4 \pm 1.7$ | $0.72 \pm 0.23$ | $1.89 \pm 0.03$ | $27.3 \pm 1.1$ |

<sup>a</sup> Pectoralis major muscle mass was lower in right vs. left limb, likely due to sham surgery in previous study.

|  | forearm amputation |  |  |  |  |  |
| --- | --- | --- | --- | --- | --- | --- |
|  | left (unaffected) |  |  | right (affected) |  |  |
| | mass<br>(g) | sarcomere length<br>(μm) | optimal length<br>(mm) | mass<br>(g) | sarcomere length<br>(μm) | optimal length<br>(mm) |
| pectoralis major | $0.27 \pm 0.15$ | $2.07 \pm 0.08$ | $31.5 \pm 10.1$ | $0.26 \pm 0.07$ | $2.21 \pm 0.07$ | $23.4 \pm 2.5$ |
| acromiodeltoid | $0.15 \pm 0.03$ | $2.18 \pm 0.03$ | $15.4 \pm 0.8$ | $0.12 \pm 0.02^*$ | $2.23 \pm 0.02$ | $11.2 \pm 0.7^*$ |
| spinodeltoid | $0.13 \pm 0.03$ | $2.19 \pm 0.01$ | $23.7 \pm 2.7$ | $0.13 \pm 0.01$ | $2.14 \pm 0.08$ | $16.8 \pm 6.9$ |
| biceps long head | $0.11 \pm 0.04$ | $2.06 \pm 0.08^*$ | $17.5 \pm 1.0$ | $0.05 \pm 0.03^*$ | $2.43 \pm 0.07$ | $10.6 \pm 1.1^*$ |
| biceps short head | $0.02 \pm 0.01$ | $1.99 \pm 0.01$ | $19.1 \pm 0.2$ | $0.03 \pm 0.01$ | $2.22 \pm 0.13$ | $15.4 \pm 2.2$ |
| subscapularis | $0.20 \pm 0.10$ | $2.14 \pm 0.09$ | $23.9 \pm 2.8$ | $0.28 \pm 0.06$ | $2.10 \pm 0.07$ | $22.3 \pm 0.9$ |
| supraspinatus | $0.24 \pm 0.11$ | $2.33 \pm 0.11$ | $24.2 \pm 1.9$ | $0.14 \pm 0.05$ | $2.31 \pm 0.25$ | $21.6 \pm 3.5$ |
| infraspinatus | $0.22 \pm 0.03$ | $2.37 \pm 0.13$ | $25.0 \pm 2.8$ | $0.17 \pm 0.03$ | $2.30 \pm 0.05$ | $21.1 \pm 1.7$ |
| teres major | $0.23 \pm 0.04$ | $2.05 \pm 0.05$ | $24.9 \pm 1.3$ | $0.22 \pm 0.06$ | $2.16 \pm 0.06$ | $22.6 \pm 0.7$ |
| triceps long head | $0.71 \pm 0.18$ | $2.03 \pm 0.05$ | $24.6 \pm 2.7$ | $0.31 \pm 0.11^*$ | $2.07 \pm 0.10$ | $17.6 \pm 2.4$ |

\*p < 0.05 for smaller value vs. contralateral limb.

In the forearm amputation group, sarcomere length was not different between limbs for most muscles. Sarcomeres were significantly longer in the affected biceps long head (17.6 ± 1.3%, p = 0.0005) compared to unaffected muscles (Table 1). However, on average optimal muscle lengths were an average of 22.7% shorter in muscles of the affected limb compared to the unaffected limb. Optimal muscle lengths were significantly shorter in affected acromiodeltoid (27.1 ± 3.9%, p = 0.010) and biceps long head (39.6 ± 3.1%, p = 0.0002) compared to the unaffected side.

Collagen content, indicative of muscle fibrosis, did not differ significantly between left and right limbs in any group.

### Group Differences

Muscle mass differed significantly across groups for biceps long head and triceps long head (Fig. 3, Table 1). Compared to the control group, the forearm amputation group had significantly lower average muscle mass in the biceps long head (51.0%, p = 0.0202) and triceps long head (57.7%, p = 0.0229). Similarly, compared to the suspension group, the amputation group had lower average muscle mass in the biceps long head (51.6%, p=0.0202) and triceps long head (56.9%, p = 0.0437). No significant differences in muscle mass were found between the forelimb suspension and control groups.

**Figure 3.**
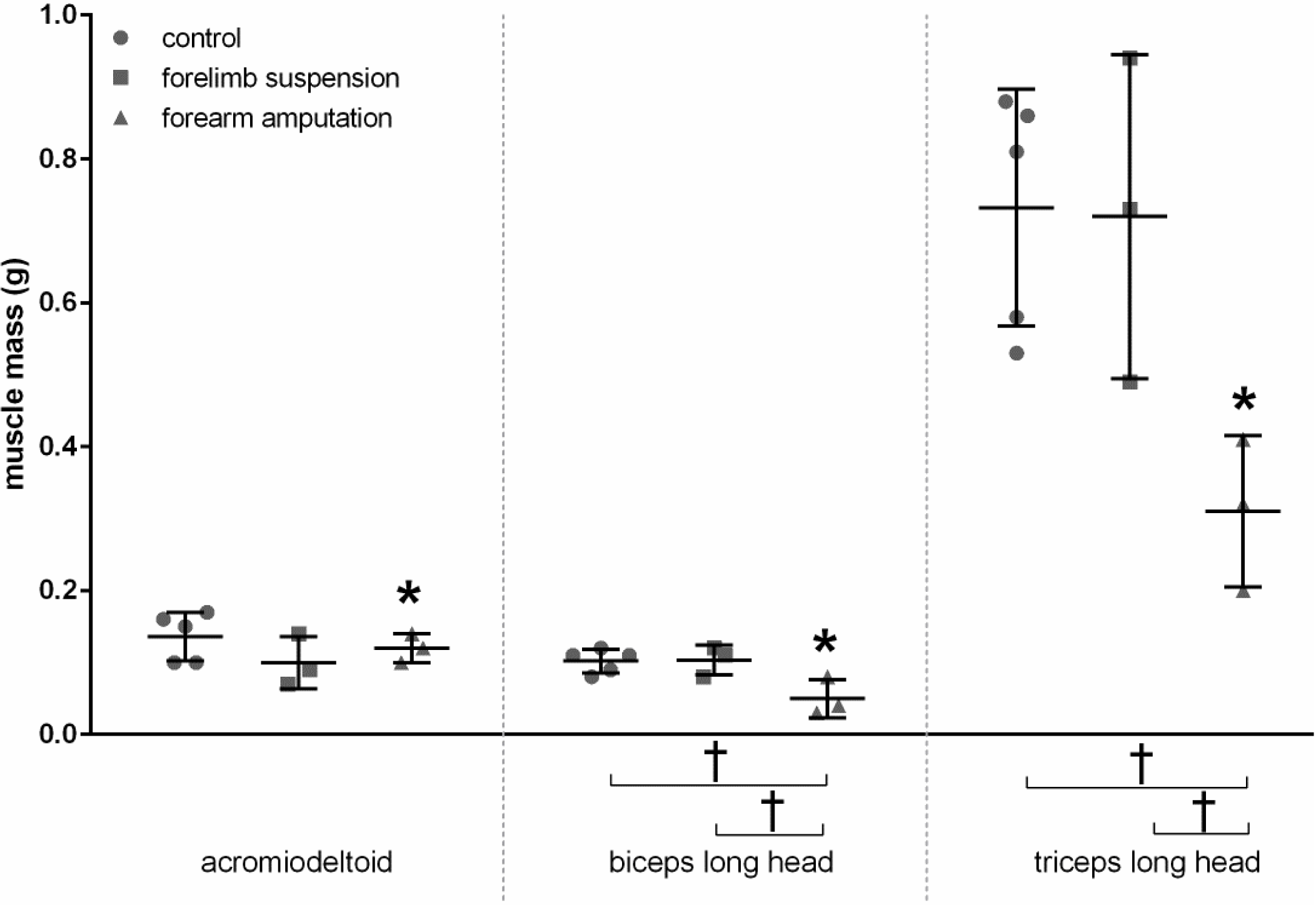
Muscle mass of forelimb muscles showing significant side-to-side differences (forearm amputation only) and group differences (control, forelimb suspension, forearm amputation). Mean ± standard deviation. *p < 0.05 for right vs. left limb. #p < 0.1 (trend) for right vs. left limb. †p < 0.05 for group comparisons.

Group differences were also observed in sarcomere length for acromiodeltoid, spinodeltoid, biceps long head, subscapularis, supraspinatus, and teres major (Fig. 4, Table 1). Compared to the control group, the amputation group had, on average, significantly shorter sarcomeres in the acromiodeltoid (10.9%, p = 0.0235) and subscapularis (10.4%, p = 0.0377) muscles but longer sarcomeres in spinodeltoid (12.6%, p = 0.000612) and teres major (25.7%, p = 0.0000366). Compared to those in the suspension group, average sarcomere length in the amputation group was significantly longer in biceps long head (17.4%, p=0.000300) and teres major (9.2%, p = 0.00212). Compared to control, the suspension group had significantly shorter sarcomeres in the biceps long head (8.5%, p=0.0449), subscapularis (13.5%, p = 0.0102), and supraspinatus (14.6%, p = 0.0401) muscles but longer sarcomeres in spinodeltoid (8.3%, p = 0.00792) and teres major (15.1%, p = 0.00148).

**Figure 4.**
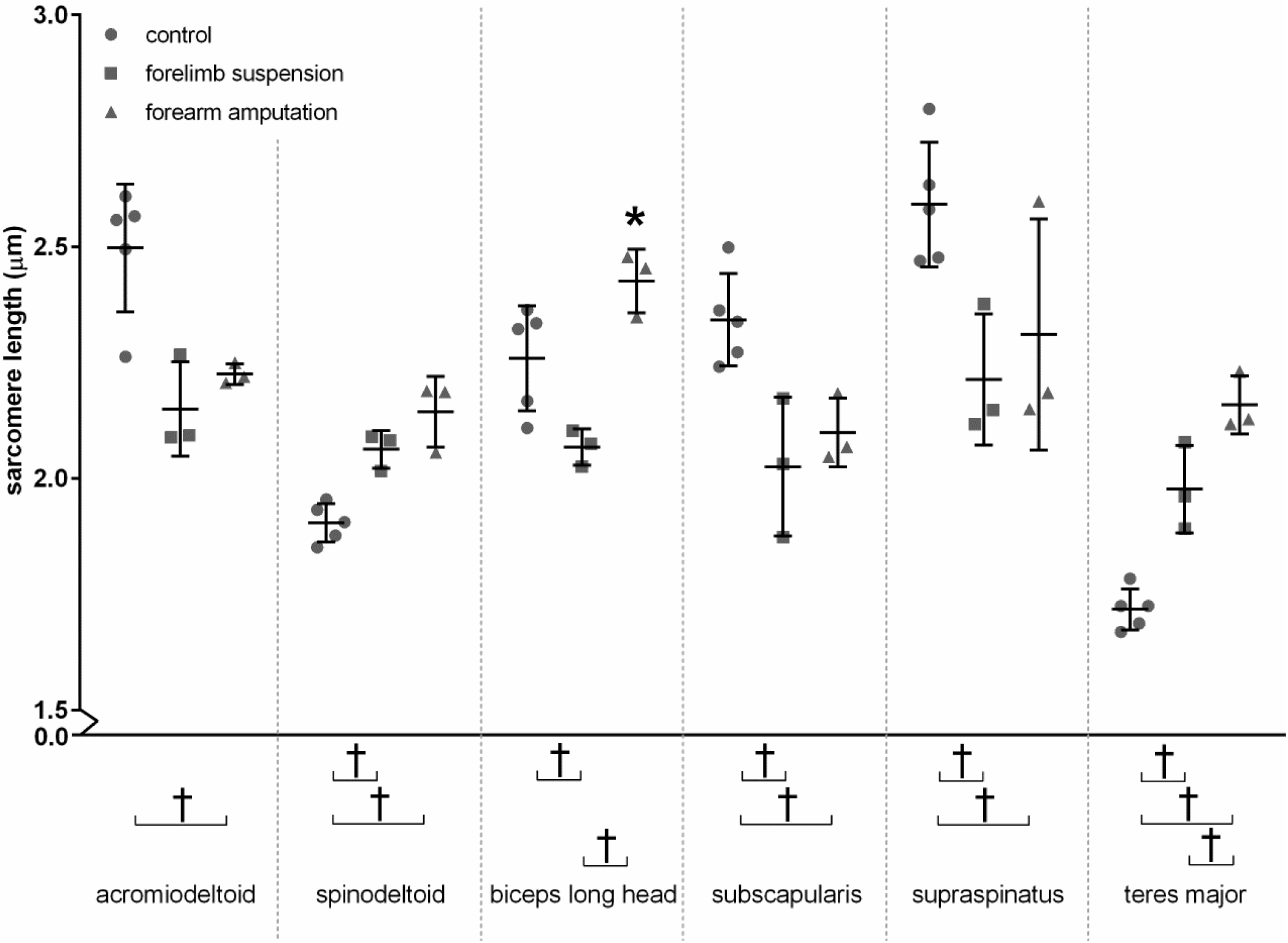
Muscle sarcomere lengths showing significant side-to-side differences (forearm amputation only) and group differences (control, forelimb suspension, forearm amputation). Mean ± standard deviation. *p < 0.05 for right vs. left limb. #p < 0.1 (trend) for right vs. left limb. †p < 0.05 for group comparisons. ‡p < 0.1 (trend) for group comparisons.

Optimal muscle length differed by group for biceps long head, biceps short head, and triceps long head (Fig. 5, Table 1). Compared to control, average optimal muscle length in the amputation group was significantly shorter for biceps long head (30.1%, p = 0.0145) and triceps long head (28.1%, p = 0.00185), indicating reduced longitudinal growth. Similarly, compared to suspension, average optimal muscle lengths in the amputation group were significantly shorter for biceps long head (36.9%, p = 0.00493), biceps short head (27.0%, p = 0.0273), and triceps long head (35.8%, p=0.000372).

**Figure 5.**
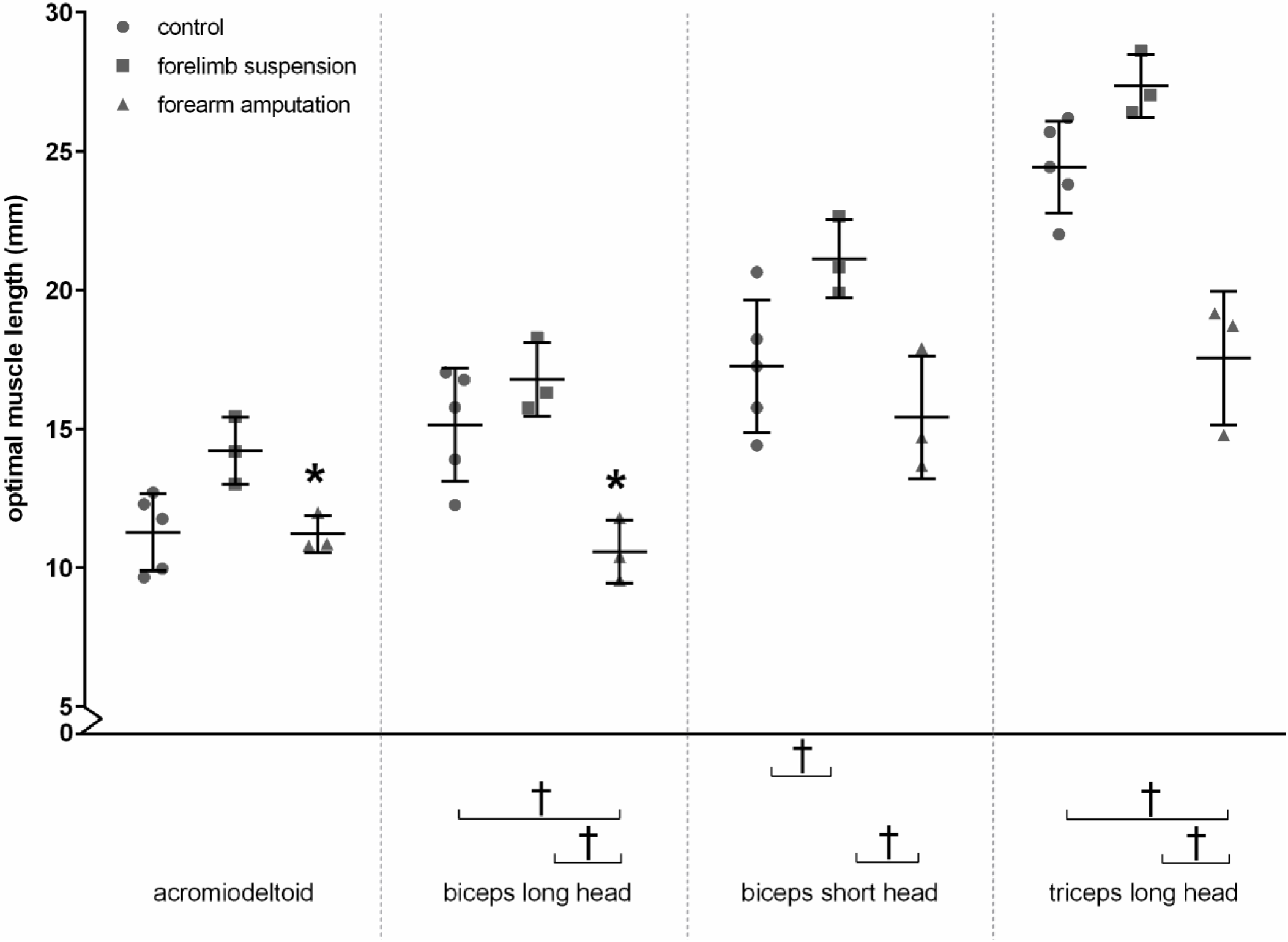
Optimal muscle lengths showing significant side-to-side differences (forearm amputation only) and group differences (control, forelimb suspension, forearm amputation). Mean ± standard deviation. *p < 0.05 for right vs. left limb. #p < 0.1 (trend) for right vs. left limb. †p < 0.05 for group comparisons. ‡p < 0.1 (trend) for group comparisons.

Qualitative analysis of histologic images revealed minimal differences across the groups in collagen staining for the 4 analyzed muscles (biceps long head, biceps short head, upper and lower subscapularis muscles) (Fig. 6). Quantitative analysis of these images showed that the ratio of collagen area to total muscle area did not differ significantly across the three groups for any of the 4 muscles examined (Table 2).

**Table 2.** Percent collagen content for the control (left forelimb), forelimb suspension (right forelimb), and forearm amputation (right forelimb) groups. Mean ± standard deviation.

|  | **control** | **forelimb suspension** | **forearm amputation** |
| --- | --- | --- | --- |
| biceps long head | 5.9 ± 1.3 | 5.9 ± 0.8<sup>a</sup> | 5.9 ± 0.0<sup>b</sup> |
| biceps short head | 6.0 ± 1.3 | 4.7 ± 0.3<sup>a</sup> | 7.9 ± 0.0<sup>b</sup> |
| upper subscapularis | 5.8 ± 2.0 | 4.1 ± 0.5 | 5.5 ± 1.7 |
| lower subscapularis | 8.1 ± 2.0<sup>a</sup> | 4.3 ± 1.1 | 4.4 ± 1.0 |

<sup>a</sup>Not all specimens could to be imaged.
<sup>b</sup>Not all specimens could be sectioned.

**Figure 6.**
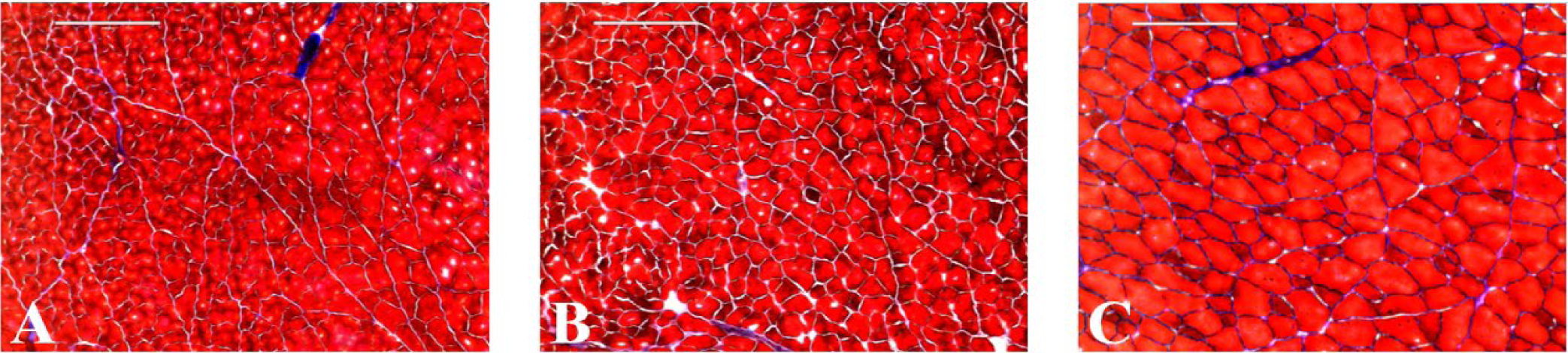
Longitudinal section of biceps long head muscle in A) control, B) forelimb suspension, and C) forearm amputation groups, stained to assess collagen area (blue) as a percentage of muscle tissue area (red). Scale bars = 200 µm.

## Discussion

Unloading with the two models had different effects on the growth of forelimb muscles. The suspension group did not affect muscle mass oroptimal muscle lengths relative to control, except for an increased optimal length for biceps short head. In contrast, the amputation intervention led to lower muscle mass and optimal muscle length for several muscles in the affected forelimb compared to the contralateral limb, suggesting that unloading via forearm amputation during postnatal development can inhibit muscle growth. Specifically, muscle mass was lower in the acromiodeltoid, biceps long head, and triceps long head, and optimal muscle length was shorter in the acromiodeltoid and biceps long head following amputation. The amputated group also had lower muscle mass and shorter optimal muscle length in the affected limb compared to the forelimb suspension (right affected limb) and control (unaffected limb) groups. On average, the amputated biceps and triceps long head muscles were approximately half the mass and 75% of the optimal length of the corresponding muscles in the suspended and control groups.

The amputation procedure provides an explanation for the specific affected muscles in this group. The biceps long head, biceps short head, and triceps long head originate at the scapula and insert to the proximal radius or ulna. During the amputation procedure, the severing at the insertion point releases the muscles and causes widespread denervation and atrophy, leading to reduced muscle mass^52^. In other studies found that denervated extensor digitorum longus muscle mass in growing rats increased after initial atrophy, following similar growth patterns as the control contralateral limbs, but soleus muscle mass decreased relative to the control^53^. The authors suggested that the increased growth was due to elevated protein synthesis after continued lengthening of the muscle, while the decrease in growth was attributed to a reduction in protein synthesis after continued shortening of the muscle. Although the biceps short head was denervated, it likely experienced extended periods of lengthening, causing it to grow similar to the suspension and control groups. The biceps and triceps likely experienced shortening over the duration of the study due to the release at amputation, which contributed to muscle mass loss and shortening. The other forelimb muscles were not affected by the amputation procedure and therefore there was no marked differences in muscle architecture.

The suspension group also exhibited changes in muscle architecture relative to the control group, which may be explained by the relative immobilization of the limbs. For example, immobilization in innervated lower limb muscles in growing rats found that a decrease in muscle mass compared to a control was attributed to higher levels of protein breakdown and reduced protein synthesis in the affected muscles when the muscles were held in a shortened position^54^. When held in a lengthened position, immobilized muscles in the lower limbs of growing rats exhibited slightly increased muscle mass compared to the control, which was attributed to decreased protein breakdown and increased protein synthesis during active and passive activity of the muscles^54^. The rats in the suspension group, although immobilized in the upper limbs, experienced typical muscle activity as seen in control rats while performing daily eating and cleaning activities, so muscle mass was not significantly affected by limb unloading because rats had full mobility of the unloaded limbs.

Altered loading had a broader and more varied impact on sarcomere length. Within the amputation group, the biceps long head had shortened optimal length but longer sarcomere length on the affected side compared to the unaffected side. The suspension and amputation groups exhibited similar changes in muscle sarcomere length across the different muscles, with shorter sarcomeres in subscapularis, and longer sarcomeres in spinodeltoid and teres major, compared to control, with additional varied effects in the acromiodeltoid and biceps long head. Hindlimb unloading has been shown to reduce titin density in the adult female rat soleus and plantaris muscles^55^. Since titin plays an integral part in sarcomere positional stability, significant losses in titin composition causes vast changes in contractile activity and likely force production. Sarcomere length contributes to optimal muscle length and muscle force production. Any deviation from the optimal sarcomere length causes actin and myosin to inefficiently interact, which limits force production in the muscle. Force production in a muscle-tendon unit, however, is not only governed by muscle length; cross-sectional area is directly proportional to the amount of force each muscle can harvest^49^ and is likely affected by changes in muscle mass^51^. Although sarcomere lengths for the anteriodeltoid, spinodeltoid, subscapularis, supraspinatus, and teres major were markedly different in the unloading groups, optimal muscle lengths for these muscles remained the same compared to the control group. Since optimal muscle length is a ratio of sarcomere length to measured muscle length, the unchanged optimal muscle length across groups is likely due to similar changes in sarcomere and optimal muscle lengths. Based on this, the biceps long head and triceps long head, which displayed remarkably lower muscle mass and longer sarcomere lengths that translated to shorter optimal muscle lengths, experienced the greatest decrease in muscle-tendon force production across the board.

These results are consistent with a previous study that investigated changes in muscle architecture in growing rats after neonatal injury to the brachial plexus nerve^51^. When comparing the affected limb to the contralateral limb, muscle mass in the same ten muscles as in this study was significantly less in all but one observed muscle, including the biceps long head and triceps long head, similar to the amputation group in this study. Concurrent to the previous study, sarcomeres in the amputation group were significantly longer in the teres major and biceps long head, along with the teres major in the suspension group relative to the control group. Unlike the injury groups, the suspension group, however, did exhibit shorter sarcomeres in the biceps long head compared to the control group. This comparison shows that muscle mass and sarcomere length in the amputation group more closely mimicked those of injury groups seen in the literature. Optimal muscle length was shorter in the biceps long head for both unloading groups, and biceps short head for the suspension group, which showed that the suspension group more closely resembled the injury groups seen in literature. The triceps long head was significantly affected by both unloading methods, but not by nerve injury, which could mean that the triceps long head muscle length is more sensitive to changes in loading than denervation.

Altered loading did not have a significant impact on muscle collagen content, with similar amounts of fibrosis between limbs, as well as across the amputation, suspension, and control groups. These findings are consistent with previous studies in adult female rat soleus muscle after 2 weeks of hindlimb unloading via tail-casting^11^. Although muscle fibrosis has been observed in children with neuromuscular disorders like cerebral palsy^42^ and with nerve injury^43,51^, our results indicate that fibrosis is unlikely to result as a consequence of reduced loading, and may instead result from other factors such as direct tissue injury or other physiological consequence..

Results from previous unloading studies vary, depending on animal age and method of unloading. Aprevious study investigating the effects of zero gravity on muscle in 3-month old mice found that mass in three leg muscles were not significantly affected by 30 days in space where observed grooming rate was high^9^. Since the mice maintained daily grooming activity, the muscles were activated throughout unloading, and these results are similar to our forelimb unloading condition with limited muscle effects. Previous hindlimb unloading studies reported muscle atrophy and decreased muscle mass. One study examined the effect of 30-day space flight on 19-week old mice and found that hindlimb muscle mass was not significantly affected by weightlessness but trended towards decreased soleus and extensor digitorum longus mass in the unloading groups^9^. This could be close to the cut-off of growing and adult. Another study using a tail-casting hindlimb unloading model in young adult female rats found that the addition of combined isometric, concentric, and eccentric muscle stimulation dampened muscle mass loss compared to the untrained contralateral limb, and muscle mass was unchanged from the regular weight-bearing group^44^. This could help adult rats maintain their muscle mass during unloading. In a partial weight-bearing study, 10-week old adult female mice gastrocnemius muscle was found to be significantly lower mass than that of the control groups^36^. Another hindlimb unloading study with adult male rats found that soleus, plantaris, adductor longus, gastrocnemius, and tibialis anterior muscle mass was significantly reduced after hindlimb unloading compared to typical weight-bearing. Isometric exercise attenuated the effects of unloading in the soleus by 54%. Isometric exercise, however, did not aid the gastrocnemius and plantaris in maintaining muscle mass, as they were significantly less than control by 15%^45^. Hindlimb unweighting was further determined as a cause for reduced muscle mass in adult rats^47^. Soleus muscle mass was significantly reduced in hindlimb unloaded growing rats compared to control rats after 17 days of unloading^46^. This effect was reversed after a 28-day reambulation period. The authors noted that during hindlimb unloading, the ankle was plantarflexed, which caused shortening of the soleus and reduced muscle mass^54^. The mechanism of unloading largely affects muscle properties. If the limb is held in place by a cast, it could be immobilized in a shortened position, which has been shown to have detrimental effects on muscle. In a model in which the unloaded limbs are exposed, they can be held at a natural, optimal position, which may not have as much of an effect of muscle.

Our forelimb suspension system differs from hindlimb suspension systems in that, although the suspended limbs are non-weight bearing, they still experience some non-weight-bearing loading and muscle activity during daily grooming and eating activities. Because muscle mass and optimal muscle length were, for the most part, similar between the suspension and control groups, this small amount of loading seems sufficient to stimulate normal forelimb muscle growth. However, our forearm amputation group experienced both reduced weight bearing and reduced limb use following amputation and were unable to walk on or groom with the amputated limb normally. The affected limb served only as an occasional weight-bearing stabilizer, and forelimb muscle use during these daily activities was greatly reduced. Therefore, the reduced muscle mass observed in this group, compared both to the contralateral limb and to the suspension and control groups, may result from limb disuse rather than direct unloading of the muscles.

### Limitations

Both male and female rats were compared together, and sex differences were not considered. Young (3-month old) male rats have shown to display greater reduction in total body mass compared to control rats over the hindlimb unloading period, while females did not^48^. The amputation group comprised of 3 male rats, whereas the other groups had at least one female, which could help explain why the amputation group displayed greater significance. The forelimb suspension group was sacrificed one week later than the forelimb amputation and control groups to accommodate the four male rats that were removed from the suspension system for a brief 4-day period due to elevated stress, as indicated by lesions underneath the harness and porphyrin discharge around the nose and eyes^36^. The removal occurred during the third week of unloading, but the rats progressed normally after the wounds healed. In the future, an additional layer of breathable fabric should be placed between the harness and rat to reduce the amount of chaffing and discomfort over the long unloading period. The suspension system, while removing weight bearing from the forelimbs, did not completely eliminate loading, as the animals were able to continue normal grooming and feeding activities, as noted above. With forearm amputation, because the affected limb experienced reduced weight bearing and overall use, the contralateral limb likely was loaded more throughout the study, potentially augmenting the side differences observed. Nevertheless, similar muscle changes were observed for the affected limb compared to the normally loaded limb of the control group.

## Conclusions

Altered loading affected upper forelimb muscle mass and optimal muscle length, primarily in the biceps and triceps muscles of the forearm amputation group. The forelimb suspension group did not experience marked differences from the control group, showing that this unloading paradigm did not negatively impact muscle growth and function, as in the forearm amputation group. Our results suggest that even limited amounts of forelimb loading during non-weight-bearing activities offset the unloading detriments observed in hindlimb unloading models, and general limb use is more important for muscle growth than weight bearing. The muscle responses in the amputation group more closely mimicked those results seen in nerve injury groups, making this a more suitable model to assess isolated muscle effects due to forelimb unloading.

